# The phased diploid genome assembly of *Vitis vinifera cv*. Shiraz

**DOI:** 10.1101/2022.12.12.520163

**Authors:** Cristobal A. Onetto, Christopher M. Ward, Anthony R. Borneman

## Abstract

Shiraz (Syrah) is a dark-skinned cultivar of the wine grape *Vitis vinifera* that forms the basis of some of the world’s most iconic wines. Worldwide, Shiraz is the fourth most planted grapevine cultivar, however it represents the most planted cultivar in Australia. Given the importance of Shiraz to worldwide wine production, a long-read based reference genome assembly for this cultivar was produced and used to investigate the unique genetic diversity and ancestral origins of this iconic variety. Phylogenetic reconstruction using high-quality genome assemblies for wine grape cultivars provided further support of a kinship between Shiraz and Pinot Noir. Harnessing long-read data, transposable element insertions potentially affecting gene function were characterized in Shiraz and assessed relative to other cultivars. This revealed a heterogenous landscape of transposon insertion points across cultivars and uncovered a specific combination of allelic variants at the *TPS24* terpene synthase locus, which when combined with appropriate environmental triggers, may allow Shiraz to produce high levels of rotundone, the aroma compound responsible for the distinctive peppery characteristics of this cultivar.

## Introduction

Shiraz (Syrah) is a dark-skinned cultivar of the wine grape *Vitis vinifera*, which is used to create some of world’s most iconic red wines. There have been many theories surrounding the history of Shiraz, including a potential origin in the city of Shiraz in ancient Persia (now a part of Iraq). However, DNA-based marker analysis has identified Shiraz as the offspring of the grape cultivars Dureza (dark-skinned) and Mondeuse Blanche (white-skinned), two cultivars that are considered native to the northern Rhône in the south-east of France [1]. It is therefore likely that Shiraz also originated in this geographic location through a natural outcrossing event, which may date back to Roman times.

Globally, Shiraz is the fourth most planted grapevine cultivar in the world. France contains the largest plantings of Shiraz, where it represents the third most common planted wine grape. In Australia, Shiraz is the most widely planted cultivar, with the 40,000 hectares positioning the country as second only to France in worldwide plantings of Shiraz. Australia is also home to many of the oldest Shiraz vineyards in the world, with vines predating the devastation of grapevine phylloxera on European vineyards in the 1800s.

One of the trademark flavours of Shiraz and especially of Shiraz grapes grown in cool-climates, is black pepper [2]. These peppery notes have been attributed to the presence of the highly-potent sesquiterpene rotundone with reported detection thresholds as low as 16⍰ng/L in red wine and 8⍰ng/L in water [3]. Although being reported in comparatively higher concentrations in Shiraz, rotundone has also been detected in several other cultivars, with Duras, Vespolina and Grüner Veltliner also displaying relatively high concentrations of this compound [4]. While the complete biosynthetic pathway of rotundone has not been elucidated, it has been shown that the sesquiterpene synthase *VvTPS24* is responsible for the synthesis of the precursor of rotundone, α-guaiene [5]. Rotundone can then be formed from α-guaiene either by simple oxidation or enzymatically through the cytochrome P450 α-guaiene 2-oxidase [6,7].

There are thousands of distinct cultivars of *V. vinifera* that are used for wine production, which display extensive phenotypic diversity. Given the economic importance of this species, genome sequencing is being used to determine the genetic differences that separate the various types of wine grapes. Early efforts in the production of reference genomes for *V. vinifera* were confounded by high levels of hetero- and hemi-zygosity [8], such that inbreeding was used to produce a homozygous line derived from Pinot Noir for initial attempts at assembling a complete grapevine genome [9]. Advances in “long-read” sequencing and phased genome assembly algorithms have now allowed for the production of highly-contiguous assemblies for the grapevine cultivars Chardonnay [10,11], Cabernet Sauvignon [12], Carménère [13], Zinfandel (syn Primativo) [14], Nebbiolo [15], Cabernet Franc [16], Riesling [17] and Merlot [18]. These studies have expanded the knowledge on the mechanisms of genome evolution in this species, highlighting the importance of structural variants and repetitive elements as drivers of cultivar and clonal phenotypic diversity.

Given the importance of Shiraz to worldwide wine production, a reference genome assembly for this cultivar is required. Long-read ONT data was selected to produce a highly contiguous diploid genome assembly for Shiraz, which could provide the basis for detailed phylogenetic investigations and to compare structural variations across Shiraz and other cultivars for which high-quality, phased genomes were available. Overall, this study aims to provide a resource for future comparative genomics of grapes with implications for diploid genome evolution, determination of clonal variation and historical provenance of *V. vinifera sativa* varieties.

## Materials and Methods

### Sampling, DNA extraction and whole genome sequencing

DNA was extracted from early season *Vv vinifera* Shiraz clone 1654 leaves taken from field grown plants at the Coombe Vineyard (Waite Campus, University of Adelaide, Adelaide, Australia). Samples were immediately frozen and ground to powder in liquid nitrogen. Approximately 100 mg of plant material was used for DNA extraction using a DNeasy Plant Mini Kit (QIAGEN, Australia) following the manufacturer’s instructions. Prior to ONT sequencing library preparation, high molecular weight DNA >10 kb was enriched using a Short Read Elimination kit SRE XS (Circulomics, Australia). Sequencing libraries were prepared using the SQK-LSK110 kit and loaded into two FLO-MIN106 and one FLO-MIN111 flow cells. Fast5 files were base-called using Guppy v. 5.0.16 (Oxford Nanopore Technologies, Oxford, UK) with the ‘sup’ model and a minimum quality score filtering of 7. A total sequencing yield of 30,663 Mb was obtained (63-fold coverage) with an N50 length of 21.8 kb. For short-read sequencing, genomic libraries were prepared using the Illumina DNA Prep library kit and sequenced on a Novaseq 6000 instrument using an S4 flow cell and 2×150bp chemistry (Ramaciotti Centre for Genomics, NSW, Australia).

### Genome assembly and annotation

Preliminary assemblies were performed using Canu v. 2.1.1 [19] and Flye v. 2.8.3 [20] and then polished with ONT reads using Medaka v. 1.5.0 (https://github.com/nanoporetech/medaka). Both assemblies were then combined using quickmerge v. 0.3 [21] with a minimum overlap of 20 kb and polished twice using short-reads and Pilon v. 1.24 [22]. Lastly, allelic contig reassignment was performed using Purge Haplotigs v. 1.1.2 [23] and assessed with BUSCO v. 5.3.2 [24] using the embryophyta ODB v10 database. A scaffolded version of the primary assembly was created for visualization purposes using the *V. vinifera* reference genome (accession GCA_000003745.2) and RagTag v. 2.1.0 [25].

A custom repeat library was built for Shiraz using RepeatModeler v. 2.0.3 [26] and the LTR pipeline extension, which applies LtrHarvest and Ltr_retriever [27] during *de novo* repeat identification. Identification of miniature inverted-repeat transposable elements (MITEs) was performed using MITE-Tracker [28]. The custom repeat sequences were combined into a single library and used for repeat annotation using RepeatMasker v. 4.1.2 [29]. Gene prediction was performed following the funannotate pipeline v. 1.8.13 [30] including Genemark-ES v. 4.68 [31], SNAP [32], Augustus v. 3.3.3 [33] and Glimmerhmm v. 3.0.4 [34] annotations allowing a maximum intron length of 10 kb. Previously published RNA-seq data for Shiraz ([35], Table S1) and the protein data of the *V. vinifera* reference genome (accession GCA_000003745.2) were provided as evidence for gene model prediction.

Homo and hemizygous regions were investigated by mapping short read data to the primary assembly. Heterozygous SNPs were called using VarScanv2.3 [36] and read-depth and SNP density calculated in 50 Kb windows (25 Kb steps) using BEDTools 2.30.0 [37].

### Phylogenetics and Identity By Descent (IBD)

Single Copy orthologs (SCO) were identified in Nebbiolo, Chardonnay, Carménère, Zinfandel, Cabernet Sauvignon, Cabernet Franc, Riesling and Merlot along with *V. vinifera sylvestris* and *V. rotundifolia* using the BUSCO eudicot dataset. Alignment was then carried out using MUSCLE [38]. To ensure that errors in annotation do not bias phylogenetic reconstruction, each alignment was manually investigated and trimmed to remove mis-annotated exons between transcripts. Each gene alignment was imported into BEAST2 [39] with unlinked Hasegawa Kishino Yano site and relaxed log normal clock [40] model priors. A MCMC chain was then run across 1×10^8^ samples with a yule tree prior. Tracer was used to identify when the model began to mix and select an appropriate burn-in (48%).

Previously published RAD-Seq lllumina paired-end reads from cultivars Dureza and Mondeuse Blanche [41] were mapped to the Shiraz primary assembly using Minimap2 v. 2.24 [42]. Mapped reads had their variants called (MQ>20) and were filtered (DP>5, GQ>20, F_MISS=0) using BCFtools v1.16 [43]. Heterozygous sites in Shiraz were further filtered to remove any sites where allele depth did not fit a binomial distribution, thereby removing somatic mutations developed after the cross event between Dureza and Mondeuse blanche. Putative alleles that suffered from dropout due to mutations in the RADtag cut site were identified by inspecting the first four base pairs of both the forward and reverse reads for mutations within Shiraz with geaR v0.1 [44]. Any tags that contained mutations were removed from the analysis space. Filtered variants were converted to the GDS format and IBD calculated using SNPrelate v1.32 [45].

### Characterization of TE content and comparative genomics

Read depth and structural variant information was leveraged using plyranges v1.18 [46] and geaR v0.1 [44] to collapse fragmented transposable element annotations into a single record. First, read depth was calculated against the Shiraz primary assembly across the middle 10 bp of a TE annotation using SAMtools v1.16.1 [43]. This was then compared to the median read depth observed across surrounding coding regions and overlapped with structural variants called from long read nanopore data using Sniffles v2.0.2 [47] to determine zygosity. Homozygous annotations were then conditionally merged if adjacent annotations from the same class were both contained within the same read and were also homozygous. Heterozygous annotations were then compared to overlapping heterozygous structural variants Adjacent transposable element annotations that were both heterozygous and contained within a single structural variant were merged into one record.

Transposable element annotations within introns and 1 kb of an orthologs TSS or stop codon were extracted and intersected to identify putative genic TE insertions. Exonic insertions points were identified by first extracting gene and CDS features from the Pinot Noir reference [9] genome and annotation sourced from Ensemble Plants (PN40024.v4).

Putative genic TEs were then leveraged against Nebbiolo, Chardonnay, Carménère, Zinfandel, Cabernet Sauvignon, Riesling and Merlot long read data (see Table S1 accession number) structural variants using Sniffles v2.0.2 [47].

Publicly available short read data from 24 common cultivars (see Table S1 for accession number) was used to determine if the TE insertion in *VvTPS24* is present in other varieties. Data was first inspected for quality using ngsReports v2.0.1 [48], mapped to the Shiraz primary assembly using BWA-mem v0.7.17 [49] and filtered to remove any reads with MQ below 20 using SAMtools v1.16.1 [43]. Both the upstream and downstream breakpoints of the TE insertion were manually inspected for reads that contained both the *VvTPS24* coding region and TE sequence and for paired reads whose insert spanned the TE insertion point (*i.e*. Reads that mapped to *VvTPS24* exon 5 whose mate mapped to the TE). For a sample to be considered to contain the TE, evidence must have been found at both the upstream and downstream breakpoints. The zygosity of the TE in each variety was then confirmed using sliding 31mers of each breakpoint and the reconstructed functional allele with Jellyfish v2.3 [50].

## Data availability

The sequencing data and genome assembly included in this study are publicly available at NCBI under BioProject PRJNA901650. The GFF files with the gene annotations are included in the supplementary data.

## Results and Discussion

### Haplotype phased assembly of the cultivar Shiraz

Recent advances in long-read DNA sequencing technologies have enabled the creation of high-quality, diploid genome assemblies for repeat-rich and highly heterozygous plant species such as *V. vinifera* [12]. Haplotype phased genome assemblies have been produced for a handful of the most widely planted cultivars, including Cabernet Sauvignon [12], Merlot [18] and Chardonnay [10,11], however to this date there is no publicly available reference genome for Shiraz, despite it being the fourth most planted cultivar in the world. To address this knowledge gap, a reference genome for the cultivar Shiraz was produced using a hybrid sequencing approach that included 63-fold coverage of ONT long reads and 60-fold coverage of Illumina short-reads. Clone 1654 was selected as the sourced plant material due to its widespread use in the Australian wine industry.

After assembly, polishing and haplotype phasing a 476 Mb long primary assembly with an N_50_ of 1.9 Mb was obtained, falling well within the expected haploid size for *V. vinifera* and only 2% smaller than the inbred Pinot Noir reference genome (PN40024). While larger primary assemblies have been obtained for other wine cultivars (Cabernet Sauvignon:590 Mb, Cabernet Franc: 570 Mb, Merlot: 606 Mb, Chardonnay:490 Mb, Carménère: 623Mb, Nebbiolo: 561 Mb and Zinfandel: 591 Mb), not all of the reported grapevine assemblies have been processed with tools to optimize the reassignment of allelic contigs and the larger sizes are likely due to the retention of both copies of highly heterozygous regions.

Haplotype phasing generated a total of 356 Mb of associated haplotigs with an N_50_ of 199 kb (Table 1). The primary assembly included 95.1% of BUSCO orthologues and contained 56.1% repetitive content primarily represented by gypsy and copia LTR retroelements (Table 1). After removal of repeat-associated gene models a total of 32,333 protein-coding genes were retained (Table 1).

**Table 1.** Shiraz assembly statistics.

|  | Primary assembly | Haplotigs |
| --- | --- | --- |
| Assembly size (bp) | 476,422,955 | 356,328,851 |
| Contigs | 435 | 2,408 |
| N<sub>50</sub> | 1,969,387 | 199,587 |
| Largest contig | 7,765,998 | 2,855,789 |
| Predicted proteins | 32,333 | 22,725 |
| Repetitive content (%) | 56.1 | 62.3 |
| Predicted hemizygous (bp) | 50,426,450 |  |
| Predicted hemizygous genes | 3,017 |  |
| Predicted homozygous (bp) | 55,847,349 |  |
| Complete BUSCOs | 1536 (95.1 %) |  |
| Complete and single-copy BUSCOs | 1466 (90.8 %) |  |
| Complete and duplicated BUSCOs | 70 (4.3 %) |  |

To assess the degree and distribution of hemi- and homo-zygosity across the Shiraz genome, information from read-depth and heterozygous variant density were assessed across both the primary and scaffolded assemblies (Figure 1). Selection of regions characterised by half the median read-depth and low heterozygous variant density highlighted 54 Mb (10.5%) of the Shiraz assembly as hemizygous, which is predicted to encode 3017 genes (Figure 1). Similar genome wide hemizygosity levels have been reported in distantly-related cultivars such as Cabernet Sauvignon (15.5% of genes in primary assembly) and Chardonnay (14.6% of genes in primary assembly) [11], suggesting a basal level of hemizygosity that separates pairs of parental alleles in *V. vinifera*.

**Figure 1.**
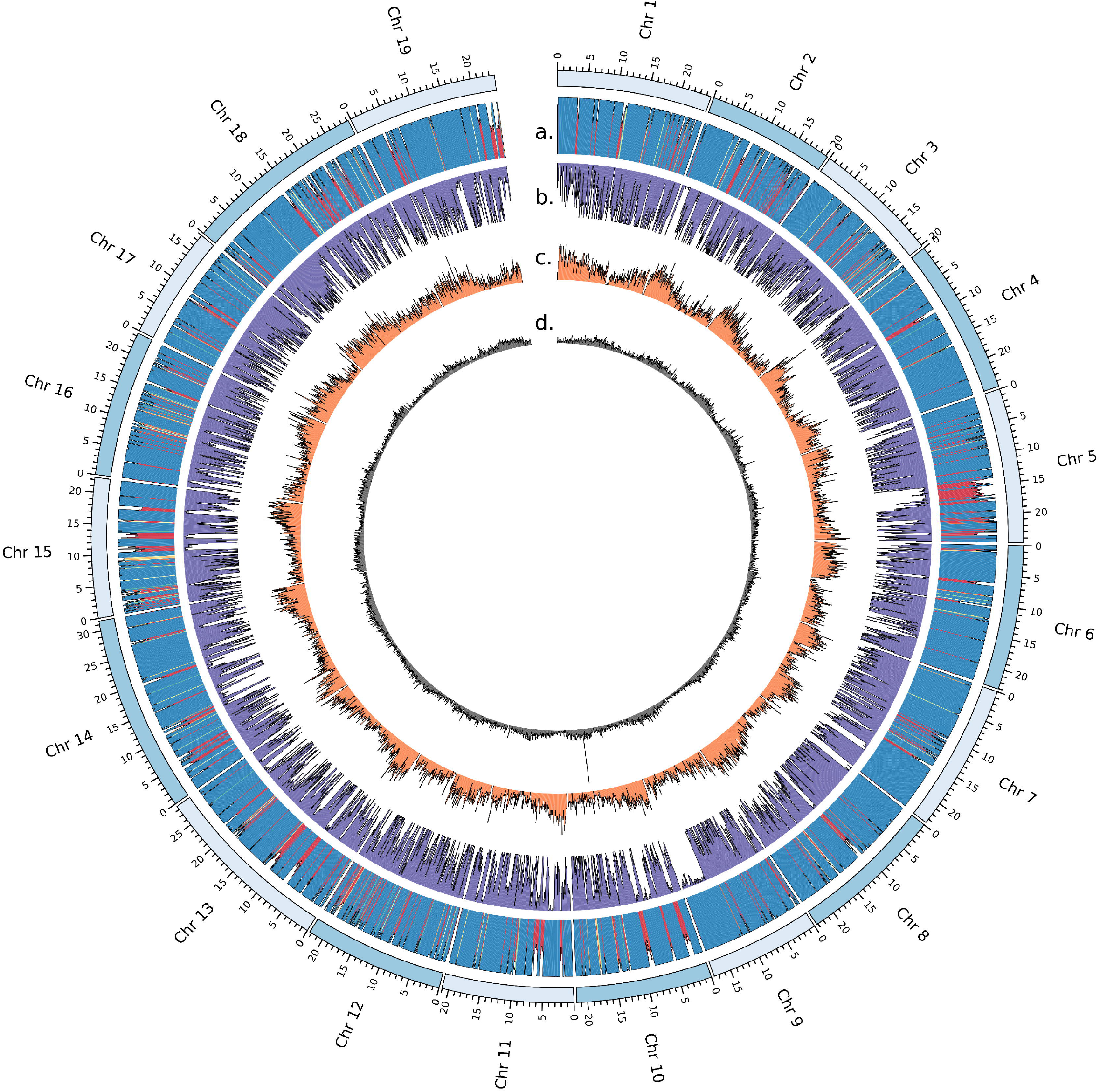
Genome assembly of *V. vinifera* Shiraz. A circos plot depicts chromosome-scaffolded primary assembly using the Pinot Noir 12X reference genome (accession GCF_000003745.3). (a) read-depth of Illumina reads mapped to the primary assembly, (b) heterozygous variants, (c) gene and (d) LTR retrotransposon density.

The functional consequences of hemizygous regions were assessed through gene ontology functional enrichment. This revealed an overrepresentation of several functional classes, including chitinases that form part of the systemic acquired resistance mechanism of *V. vinifera* [51] and terpene synthases (Table S2). This is not surprising as previous comparative genomic studies have suggested that hemizygosity and variation in gene content have a potential contribution towards the phenotypic differences between cultivars [11,15].

Homozygous regions, which were categorised as areas of median read-depth and low heterozygous variant density, comprised 11.7% of the primary assembly. The longest run of homozygosity was a stretch of 4.8 Mb located at one end of chromosome 9 (Figure 1). In comparison, Chardonnay, which has been suggested to be a naturally inbred cultivar [10], was shown to contain twice the levels of homozygosity (22.4%) as Shiraz, providing support that the parental cultivars of Shiraz do not share a recent common ancestor.

The density of genes (Figure 1c) and LTR retrotransposons (Figure 1d) displayed a distinctive pattern whereby gene density decreased, with a concomitant increase in LTR density surrounding centromeric regions. However, a clear spike in LTR density was also observed outside of the centromeric region on Chr 10 (Figure 1d), suggesting a nested transposable element insertion region. Detailed annotation of this LTR repeat domain rich region on Chr 10 identified 29 separate LTR repeat domains and 127 internal domains, which were distributed across an 88 kb region (Figure S1). Long-terminal repeat insertions appeared to be nested within the internal domains with multiple LTR domains occurring within small (<2kb) windows. Analysis of the mapped reads (Figure S2) showed no indication of a mis-assembly across the region with long reads spanning multiple nested LTR domains.

### Phylogenetic reconstruction and parentage of Shiraz

The availability of a large number of long-read grapevine genomes offered the ability to assess phylogenetic relatedness that encompassed information inherent within both phased alleles across the diploid genome. Single copy orthologs were identified across the ten long-read *V. vinifera vinifera* genomes, in addition to *V. vinifera sylvestris* and *V. rotundifolia* as outgroups. Phylogenetic reconstruction using these 437 single copy orthologs revealed Pinot Noir as the closest relative of Shiraz (Figure 2), providing further support for their proposed kinship [52]. Cabernet Franc and its offspring Cabernet Sauvignon, Carménère and Merlot (Figure 2) were closely related, yet clade topology placed Cabernet Franc as the most derived and Merlot as more closely related to Riesling than the Cabernet clade. This is likely due to alleles from the alternative parental haplotype influencing the topology of the tree and indicate that care should be taken in future studies when interpreting the topology of closely related crop plants with shared ancestry unless true haplotypes can be resolved. Despite this, the tree topology suggests Merlot and Riesling may share a close relative.

**Figure 2.**
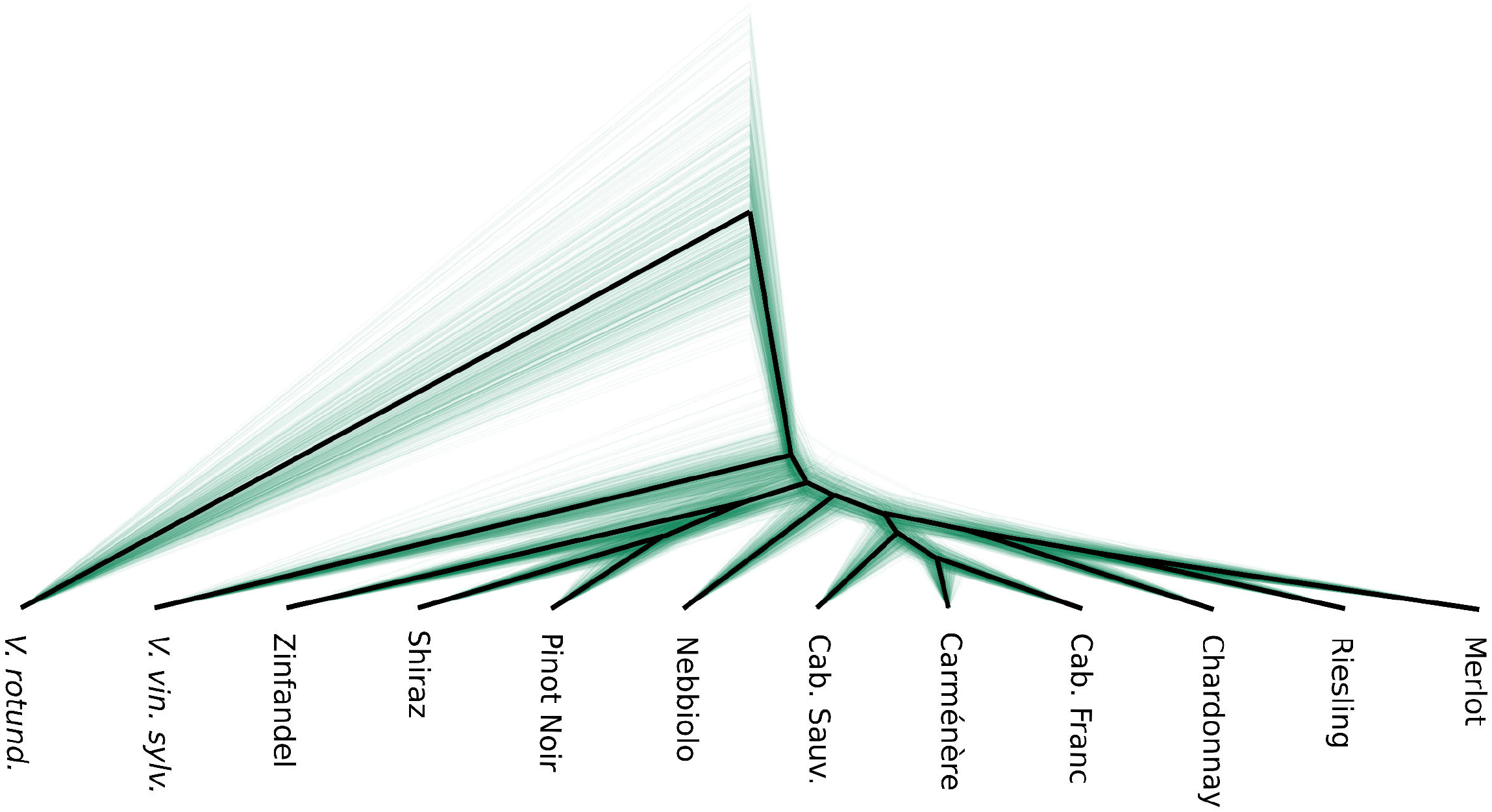
Bayesian phylogeny of Shiraz and other common varieties with *Vitis vinifera sylvestris (V. vin. sylv.)* and *Vitis rotundifolia (V. rotund.)* used as outgroups. Cab. Sauv: Cabernet Sauvignon, Cab. Franc: Cabernet Franc.

As SSR data was previously used to suggest that Mondeuse Blanche and Dureza comprise the parents of Shiraz [1], existing RAD-Seq data from these two cultivars [41] was utilized to provide further support to their relationship to Shiraz. Whole genome variant calling was performed using the RAD-Seq data of Mondeuse Blanche and Dureza against the Shiraz genome sequence, producing a space of 81,551 variants for analysis. Quality filtering, removing calls within annotated repeats and RAD allele dropout decreased the total number of usable genotypes to 22,358 for kinship estimation. During filtering, allelic ratio was used to calculate a binomial probability (np=0.5) at each heterozygous Shiraz variant to remove variants that may be somatic from the Shiraz genotypes. Kinship estimation and identity by descent (IBD) calculation was then carried out using the MOM method. A kinship matrix consistent with a parent-offspring relationship was calculated for Dureza and Shiraz (IBD=0.25, (k_0_ 0, k_1_ 1)). However, values that would be considered consistent with a true parent-offspring relationship were not recovered for Mondeuse Blanche (IBD=0.192, (k_0_ 0.23, k_1_ 0.77)). This suggests that the relationship between Mondeuse Blanche and Shiraz may be more complex than previously estimated using SSR markers.

### Repeat characterization in Shiraz reveals TEs near genes absent from other varieties

Somatic variation and specifically transposable element (TE) insertions have been shown to drive adaptation [53,54] and to provide an important source of genetic diversity for trait selection and breeding in clonally propagated crop plants [55,56]. Disruption of proper gene function via the insertion of TEs can either occur directly through disruption of the open reading frame, or indirectly through influencing transcription by insertion into regulatory regions or through chromatin availability and epigenetic silencing [57–59]. In *V. vinifera*, TE insertions have been identified as the causative mutation that underpins important phenotypes such as differences in berry color [60], which have convergently occurred across multiple lineages [60,61] through insertion of the Gret1 LTR into the promoter region of *VvMYBA1* [60,62].

Genome-wide characterisation of TE content identified 125,736 TE annotations in the Shiraz primary assembly (Figure 3a). Long terminal repeats (LTR) comprised the most frequent TE class, with Copia and Gypsy LTRs representing 50.6% of all annotated TEs. LINE elements (17.8%) were the third most common class followed by MITEs (14%) (Figure 3a).

**Figure 3.**
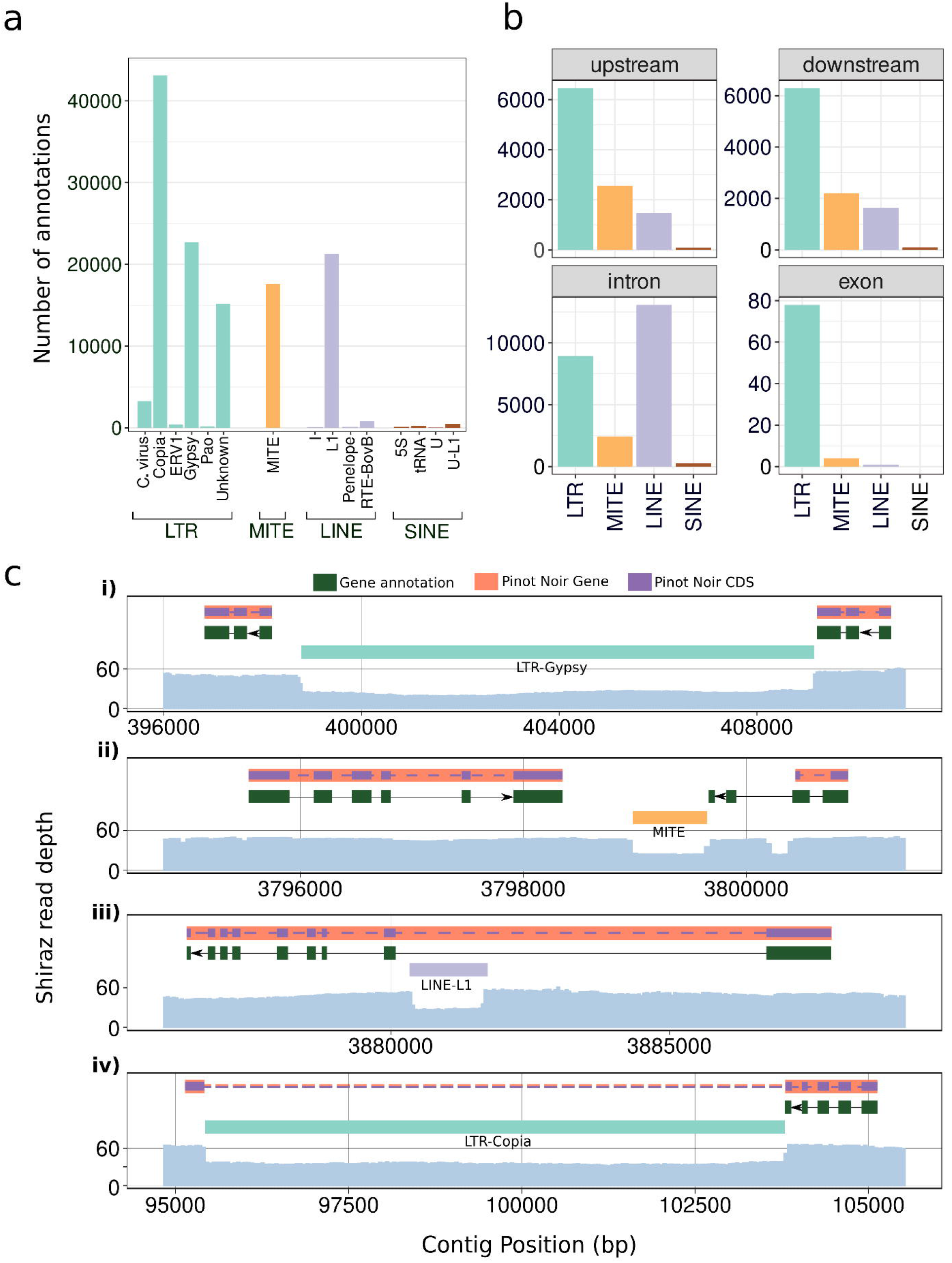
Repeat structure of *V. vinifera* Shiraz clone 1654 (a) Genome wide classification of transposable elements (TEs) annotated in the Shiraz primary assembly. LTR, MITE, LINE, RC and SINE TEs are broken down into subclasses and colored according to their class. (b) Classification of TEs that are annotated 1kb upstream, downstream and intronic of an annotated gene or exonic of a Pinot Noir gene/CDS combo. Gaps in Pinot Noir gene/CDS alignments to the Shiraz reference are depicted as dashed lines. (c) TEs that are specific to Shiraz after overlapping genic TEs with structural variants from Nebbiolo, Chardonnay, Carménère, Zinfandel, Cabernet Sauvignon, Riesling and Merlot long-read data. (i) upstream, (ii) downstream, (iii) intronic and (iv) exonic.

As TE insertions within genic regions can impact gene function these were mapped across the Shiraz genome and compared to those in other cultivars. Firstly, the Shiraz gene and repeat annotations were utilized to identify TEs within 1 kb of either the 5’ or 3’ termini of each gene model, in addition to those within intronic regions. This identified 10,570 TE annotations upstream (Table S3) and 10,223 annotations downstream (Table S3), which may affect gene regulation (Figure 3b). In contrast to the genome-wide data, in which LTR elements are the most frequent TE across the genome, MITE elements were most commonly observed upstream (24.2%) and downstream (21.6%) of genes, agreeing with past studies [57,63]. LINE insertions were the most frequently observed TE within intronic regions comprising 52.9% of the 24,721 intronic TEs (Table S3).

Insertions of TEs within exons would evade detection by the previous methodology, as it is likely that exonic insertion would interfere with correct gene annotation within Shiraz. To overcome this limitation, a secondary methodology was applied in which gene annotations were extracted from the Ensembl Pinot Noir reference genome entry and mapped to the Shiraz primary assembly. This identified a total of 31,839 putative gene annotations which were then overlapped with the Shiraz TE annotations, revealing 83 potential exonic insertions in Shiraz (Table S3) (Figure 3b).

To identify genic TE insertions that are variable between cultivars or specific to Shiraz, structural variants called from long read data of Cabernet Sauvignon, Chardonnay, Carménère, Merlot, Nebbiolo, Riesling, Shiraz and Zinfandel were cross-referenced against the set of putative genic TEs. This identified 4,554 genic TE insertions variable between these cultivars (Table S4), which may contribute to their phenotypic diversity. Shiraz-specific TE insertions were also identified (Table S5), 38 upstream (LTR: 81.6%; MITE: 10.5%; LINE: 7.9%) (Figure 3c i), 29 downstream (LTR: 93%; MITE: 7%) (Figure 3c ii), 53 intronic (LTR: 60.4%; MITE: 1.9%; LINE: 37.7%) (Figure 3c iii) and 6 exonic (LTR: 100%) (Figure 3c iv). Furthermore, the majority of Shiraz specific TEs were in the heterozygous state (85.7%).

### The *VvTPS24* locus comprises a distinct genotype in the cultivar Shiraz

From the few characterized enzymes involved in the production of aroma compounds or their precursors, terpene synthases have received particular attention due to their role in the biosynthesis of volatile terpenoids that define the varietal characters of several grape cultivars [64,65]. The potent bicyclic sesquiterpene rotundone is characterized by a peppery aroma and is responsible for one of the key varietal characteristics of Shiraz [2]. The biosynthesis of rotundone involves a two-step process, involving production of the precursor α-guaiene from farnesyl pyrophosphate by an allele of the sesquiterpene synthase *VvTPS24* [5] and the subsequent oxidation of α-guaiene into rotundone [6,7]. Wildtype *TPS24* produces an array of sesquiterpenes, of which only a minor fraction is α-guaiene [65]. However, a high α-guaiene producing variant of *VvTPS24* has been recently reported in Shiraz, which contains two polymorphisms in the active site of the protein [5]. This is likely to be linked to the ability to synthesize high levels of rotundone, although it remains to be determined if this variant is present in other cultivars.

Investigation of the *VvTPS24* locus in the diploid genome assembly of Shiraz revealed a single predicted gene model (Gene ID 002051) with 98.9% protein similarity to *VvTPS24* ortholog from Pinot Noir (NCBI accession XP_002282488) that was present in the haplotig pool. The protein predicted by this gene annotation in Shiraz contained both the T414S and V530M substitutions that have been previously associated with higher production of α-guaiene [5] (Figure S3).

Given the diploid nature of the Shiraz assembly, a second allele of *VvTPS24* would also be expected to be present in primary contigs of the genome assembly. To determine if this second allele was present, but missing from the initial annotation, splice-aware mapping of the CDS of *VvTPS24* was performed against the primary assembly. Results showed the presence of a second putative allele of *VvTPS24* that contained a large, 15 kb insertion within exon 5 (Figure 4a). Annotation of this large insertion using the RepBase database revealed the insertion to be a Ty3-gypsy–type retrotransposon with high similarity to the Gypsy18 LTR of grapevine (NCBI accession AM476928) [66]. Manual annotation of this presumably inactivated allele of *VvTPS24* indicated that, in the absence of the LTR insertion, the protein sequence of the second allele would have 99.5% similarity to the Pinot Noir *VvTPS24* and only contain the T414S substitution (Figure S3). Overall, these results indicate that due to this unique combination of SNP and structural variation in Shiraz, the main product of the VvTPS24 locus would be α-guaiene.

**Figure 4.**
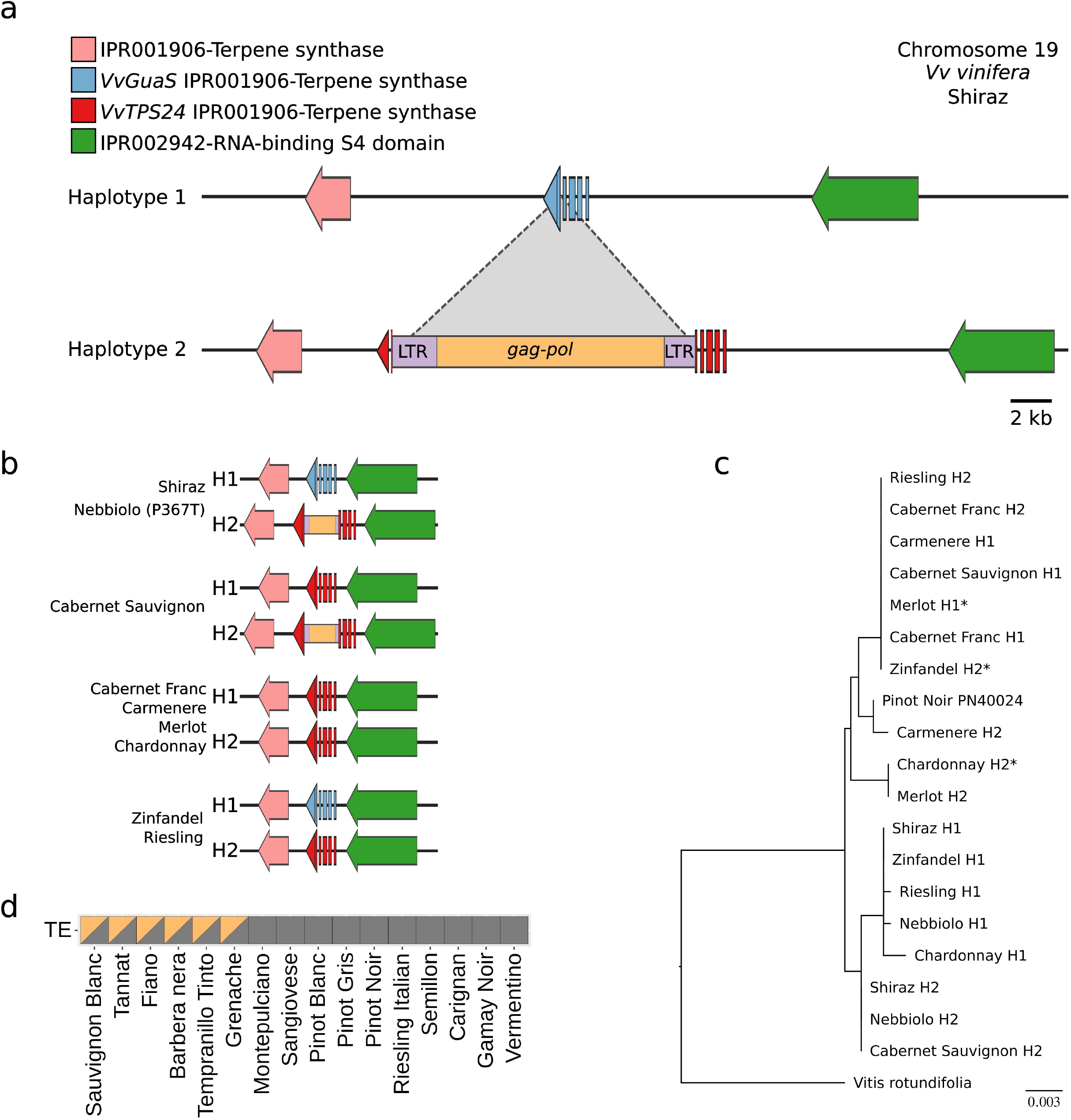
The *VvTPS24* locus in *V. vinifera*. (a) Functional annotation of the *VvTPS24* locus in the haplotypes of *V. vinifera* Shiraz. *VvGuaS* (blue) represents the polymorphic variant of *VvTPS24* containing the T414S and V530M amino acid substitutions previously reported [5]. *VvTPS24* (red) represents an allele without the T414S and V530M amino acid substitutions. Dotted lines depict the insertion point of a transposable element within exon 5 of *VvTPS24*. (b) Schematic representation of the *VvTPS24* locus in the haplotype phased assemblies of eight additional *V. vinifera* cultivars. (c) Maximum likelihood phylogenetic tree predicted from the CDS of *VvTPS24* for nine *V. vinifera* cultivars with *V. rotundifolia* as an outgroup. The CDS for all alleles including retroelement disrupted variants were manually predicted using the *VvTPS24* CDS of the Pinot Noir PN40024 reference genome. Cultivars denoted with an asterisk (*) contain an insertion/deletion within the CDS of *VvTPS24*. (d) Presence (orange) and absence (grey) of a retroelement within *VvTPS24* for 16 *V. vinifera* cultivars predicted using publicly available short-read data (Table S1).

To obtain a broader understanding of the potential metabolic pathways underpinning rotundone biosynthesis in other cultivars, the *VvTPS24, FPPS* (farnesyl diphosphate synthase) [67] and *VvSTO2* (α-guaiene 2-oxidase) [7] loci, which have all been linked to the biosynthesis of rotundone, were genotyped across the eight *V. vinifera* cultivars with available haplotype phased assemblies (Figure 4b). No structural differences were observed for either the *FPPS* or *VvSTO2* genes (data not shown), however, an allele of *VvTPS24* with a LTR retrotransposon inserted within exon 5 was observed in the cultivars Nebbiolo and Cabernet Sauvignon (Figure 4b). Alignment of the theoretical VvTPS24 proteins indicated that, like Shiraz, the *V. vinifera* cultivars Zinfandel, Riesling and Nebbiolo also carry a copy of *VvTPS24* containing the T414S and V530M substitutions (Figure 4b, Figure S3). An additional amino acid substitution (P367T) was also observed in the H1 haplotype of Nebbiolo (Figure S3). Highly variable rotundone concentrations in grapes have been linked with environmental conditions and phenological stages grape ripening [68–70], however, these results also suggest a molecular basis may prime an inherent variability of rotundone concentrations between cultivars of *V. vinifera* [4,71].

Phylogenetic reconstruction of *VvTPS24* annotations revealed that the alleles specifying the T414S and V530M amino acid substitutions share an evolutionary origin (Figure 4c). The topology of the gene tree was largely congruent with the species tree (Figure 2), with the lineages derived from Cabernet Franc being monophyletic in at least one allele. The H2 allele of Chardonnay and Merlot were also shown to be closely related (Figure 4c), providing additional support for the shared ancestry suggested by the species tree (Figure 2). After excising the TE insertion from exon 5, all alleles containing the TE were found to be monophyletic (Figure 4c), indicating that Shiraz inherited both the inactivated and high α-guaiene producing alleles of *VvTPS24* through an ancestral outcrossing event. Short-read data from a further 17 cultivars was mapped to the Shiraz primary assembly identifying the TE insertion in exon 5 of *VvTPS24* in further six cultivars (Figure 4d). While these results might suggest these cultivars share an ancestry with Shiraz, long-read sequencing data will be required to confirm the genotype of the putative functional copy of *VvTPS24* in these cultivars. Correlation of these results with measured rotundone levels in other cultivars might provide further insights into the relevance of the detected allelic differences in the *VvTPS24* locus, once the environmental triggers of sesquiterpene biosynthesis is more thoroughly understood.

## Conclusions

The availability of a reference genome for Shiraz, expands the pool of genomes available for wine grapes, while providing a foundation resource for whole-genome studies involving this iconic cultivar, including intra-cultivar variant identification and transcriptomic studies using a matching reference genome, rather than a disparate proxy. The identification of a pair of specific genomic variants involving the *TPS24* gene, outline a potential genetic basis for the propensity of Shiraz (and other cultivars) to be primed for formation of α-guaiene-type sesquiterpenes, such as rotundone, when exposed to appropriate environmental triggers. Following appropriate confirmation, this study could provide a genetic marker for production of cool-climate associated peppery characters in future grape breeding strategies.

## Supporting information

annotations.zip

## Conflict of Interest

The authors declare that they have no conflicts of interest.

## Author Contributions

C.A.O and C. M. W. performed the sequencing, data analysis and drafting of the manuscript. A.R.B supervised and provided guidance through the project and drafted the manuscript.

## Acknowledgements

This work was supported by Wine Australia, with levies from Australia’s grapegrowers and winemakers and matching funds from the Australian Government. Support for DNA sequencing was provided by Bioplatforms Australia as part of the National Collaborative Research Infrastructure Strategy, an initiative of the Australian Government. The AWRI is a member of the Wine Innovation Cluster (WIC) in Adelaide. The authors would also like to thank Benjamin Pike for providing access to the Coombe Vineyard.

## Supplementary Material

Supplementary_figures.pdf: Figures S1-S3

Tables_S1.txt: SRA accessions included in this study.

Tables_S2.txt: GO molecular function enrichment analysis of hemizygous genes predicted in the assembly of Shiraz

Tables_S3.txt: Genic transposable element insertions in the Shiraz genome

Tables_S4.txt: Variable transposable element (TE) insertions between cultivars of *V. vinifera*

Tables_S5.txt: Shiraz specific transposable element (TE) insertions.

annotations.zip: Gene annotations in GFF3 format of the Shiraz diploid assembly.

## Notes

### Competing Interest Statement

The authors have declared no competing interest.

