## Supplementary_figures.pdf for "The phased diploid genome assembly of *Vitis vinifera cv*. Shiraz"

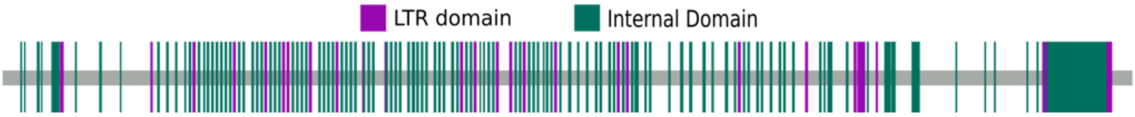


**Figure S1**. RepBase annotations for LTR region, LTR domains (initial and terminal repeats) and internal domains (ORFs and UTRs of internal proteins).

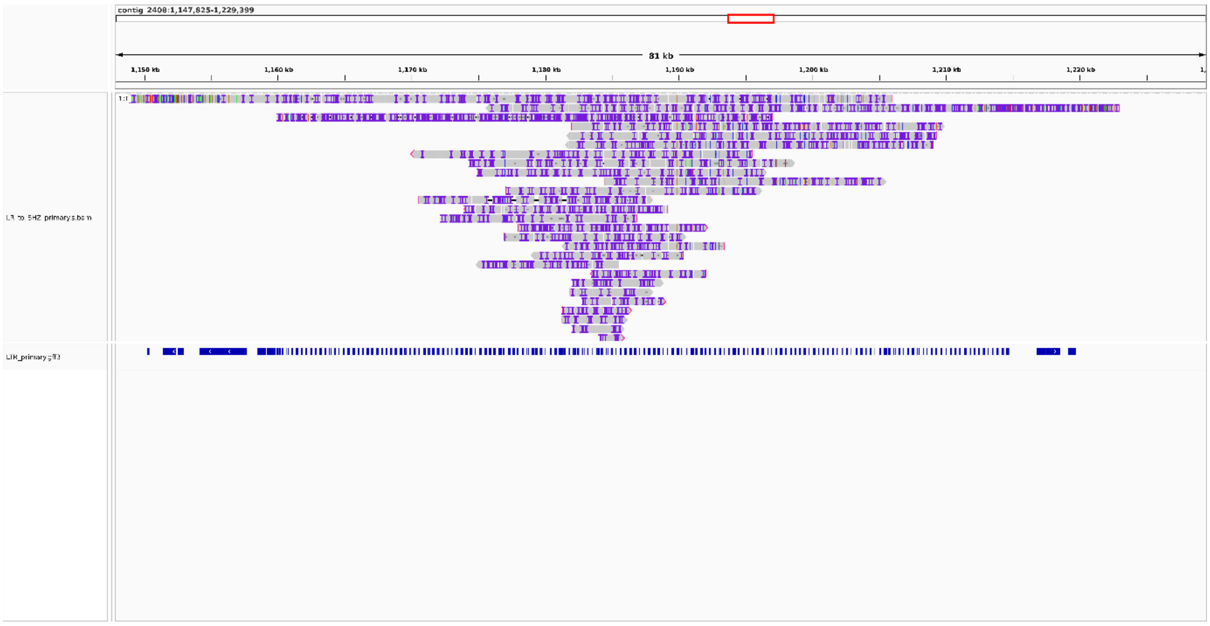


**Figure S2**. IGV screenshot of ONT reads mapped to the LTR nested region of Chr 10. First and second tracks shows mapping location of reads and repeat annotation, respectively.

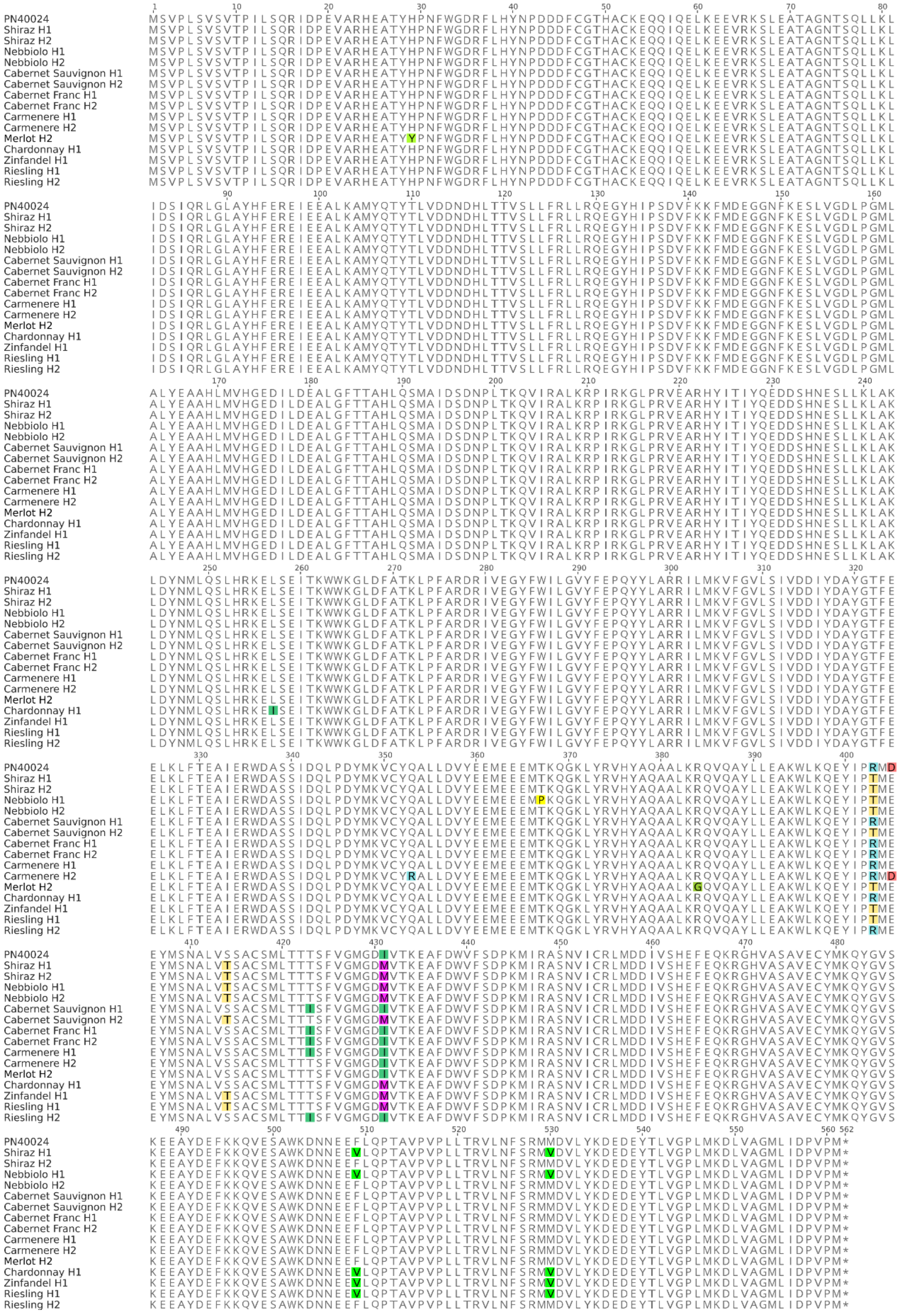

**Figure S3**. Protein alignment of predicted VvTPS24 sesquiterpene synthases in 10 *V. vinifera* cultivars. All amino acid differences between proteins are highlighted. Haplotype designation matches with locus presented in Figure 4. Only proteins predicted from CDS without detected frameshift mutations are presented.
